# Ipsilateral somatosensory cortex contains task-relevant tactile representations despite contralateral activation dominance

**DOI:** 10.1101/2025.11.13.688241

**Authors:** Hyeree Yoon, Sungbeen Park, Seojin Yoon, Sungshin Kim

**Affiliations:** Department of Data Science, Hanyang University, Seoul 04763, Republic of Korea; Department of Artificial Intelligence, Hanyang University, Seoul 04763, Republic of Korea; Department of Healthcare Digital Engineering, Hanyang University, Seoul 04763, Republic of Korea; Center for Neuroscience Imaging Research, Institute for Basic Science, Suwon 16419, Republic of Korea

## Abstract

Tactile perception has long been thought to be processed primarily in the contralateral somatosensory (S1), yet the functional role of ipsilateral S1 in human tactile discrimination remains unclear. We designed an fMRI experiment to investigate the neural representation of sequential vibrotactile discrimination as participants judged which of two successive stimuli delivered to the fingertip was higher in frequency. Contralateral S1 showed robust activation during unilateral stimulation, whereas ipsilateral S1 exhibited overall suppression. Interestingly, we found reliable frequency-dependent modulation in ipsilateral S1, which was absent in a control condition requiring no comparison between stimuli. Multivariate representational analyses further demonstrated that the difference in the neural representations between successive stimuli was strengthened under memory demands, particularly within ipsilateral S1 and parietal cortices. The ipsilateral representation was observed for stimulation of either hand, suggesting a bilateral and symmetric encoding mechanism rather than a hand-specific effect. Together, these findings demonstrate that the ipsilateral S1 contributes to tactile discrimination despite its overall suppression during unilateral stimulation, challenging classic contralateral models of somatosensory processing. Our results highlight interhemispheric coordination as a key mechanism underlying perceptual decisions.

**Significance statement:** Touch is traditionally thought to be processed predominantly in the somatosensory cortex contralateral to the stimulated body part. Here, we challenge this view using a human fMRI experiment. While the overall activity in the ipsilateral somatosensory cortex was suppressed, its multivoxel activity patterns distinguished the first and second tactile stimuli during tactile decision-making task. This representation was enhanced by mnemonic demands, appearing symmetrically for both hands. These findings reveal a dissociation between activation strength and information representation in the somatosensory cortex. Our findings suggest that tactile decisions rely on bilateral, context-dependent somatosensory representations, extending classical models that emphasize contralateral sensory processing.

## INTRODUCTION

It has been well established that tactile stimulation of one hand activates the contralateral primary somatosensory cortex (S1) and suppresses activity in the ipsilateral S1 (*1*, *2*). Early studies interpreted this asymmetry as evidence that tactile perception depends mainly on the contralateral hemisphere. However, recent work challenges this view. Ipsilateral S1 shows task-dependent modulation during tactile stimulation (*3*). Finger stimulation evokes transient deactivation in the ipsilateral S1 that is time-locked to contralateral activation due to the interhemispheric inhibition through transcallosal pathways (*2*, *4*, *5*). Ipsilateral deactivation also consistently occurs during pain stimulation, indicating that inhibitory interactions are a stable feature of somatosensory processing (*6*, *7*). Beyond inhibitory dynamics, recent work revealed that the ipsilateral S1 encodes decision outcomes during tactile learning, driven by top-down inputs from the lateral orbitofrontal cortex (*8*). This finding indicates that the ipsilateral S1 contributes to adaptive behavior by integrating feedback-related information, extending its role beyond sensory encoding. Together, these studies show that tactile perception arises from dynamic interhemispheric coordination between the two S1 cortices (*1*, *9*) and higher-order cortical feedback, which jointly shape the neural representation of touch (*8*). Yet, the distributed representation of tactile discrimination in the ipsilateral S1 remains unclear.

Indeed, most human neuroimaging studies have used univariate analyses that measure mean activation within regions, which limits insight into how information is represented. Electrophysiological studies show that neurons within the same cortical area encode different components of tactile discrimination—the first stimulus, the second, or the comparison outcome—indicating that population-level activity patterns carry task-relevant information (*10*, *11*). These population codes are not static but vary with cognitive context. When participants actively discriminate tactile stimuli rather than passively receive them, neural representations in somatosensory and parietal cortices become sharper and more distinct, reflecting task-dependent refinement of sensory coding (*12*). Consistent with this, multivariate analyses have shown that tactile representations become increasingly separable as cognitive demands rise during active discrimination or memory-based comparison (*13*). This diversity implies that spatially distributed activity patterns, rather than mean signal amplitude, capture key aspects of sensory representation.

To characterize such neural representation in the distributed activity pattern, multivariate methods such as representational similarity analysis (RSA) have been employed (*14*). RSA quantifies these representational geometries and enables direct comparison between cortical representations and behavioral performance. This approach can be applied to bilateral somatosensory regions to examine how tactile representations are organized across hemispheres and relate to perceptual accuracy. Beyond inhibitory mechanisms, the ipsilateral S1 receives top-down feedback from higher-order cortical areas, as shown in recent work (*8*), suggesting it may integrate contextual and comparison-related information to support perceptual decisions. Because feedback from such areas often carries information about stimulus comparison and choice outcomes, the ipsilateral S1 is well positioned to represent relational rather than absolute sensory codes. This integrative function could transform the ipsilateral S1 from a suppressive relay into a computational node that encodes decision-relevant differences between stimuli.

Building on these ideas, we hypothesized distinct computational roles of the contralateral and ipsilateral S1 in representing the absolute frequency of tactile input and higher-level information related to tactile decision-making. In this paper, we report a surprising finding that the perceived sequential tactile difference is primarily represented in the ipsilateral S1, rather than in the contralateral S1. These effects were more pronounced during the active discrimination task than during the passive stimulation task, reflecting the role of cognitive demands in enhancing the representation of task-relevant tactile information. Importantly, these results are not hand-specific; they were replicated across two independent experiments involving stimulation of the right and left hands, supporting a consistent role of the ipsilateral S1 in tactile discrimination.

## RESULTS

### Behavioral performance

In Experiment 1, participants reliably discriminated between vibrotactile frequencies around the 15 Hz reference (Fig. 1C). The proportion of “judged higher” responses increased monotonically with comparison frequency, showing a typical psychometric curve (*15*). Overall accuracy was 0.87 ± 0.02 with 1.50 ± 0.23 Hz (mean ± SEM) JND (Just Noticeable Difference), demonstrating participants’ robust sensitivity to frequency differences. Similarly, participants in Experiments 2 and 3 showed perceptual accuracy comparable to that in Experiment 1 (Experiment 2: 0.86 ± 0.02 with a JND of 1.62 ± 0.18 Hz, Experiment 3: 0.82 ± 0.02 with a JND of 2.07 ± 0.30 Hz). These results indicate that participants maintained high levels of engagement and discriminative performance across all experimental conditions, regardless of the stimulated hand or the response modality.

**Fig. 1.**
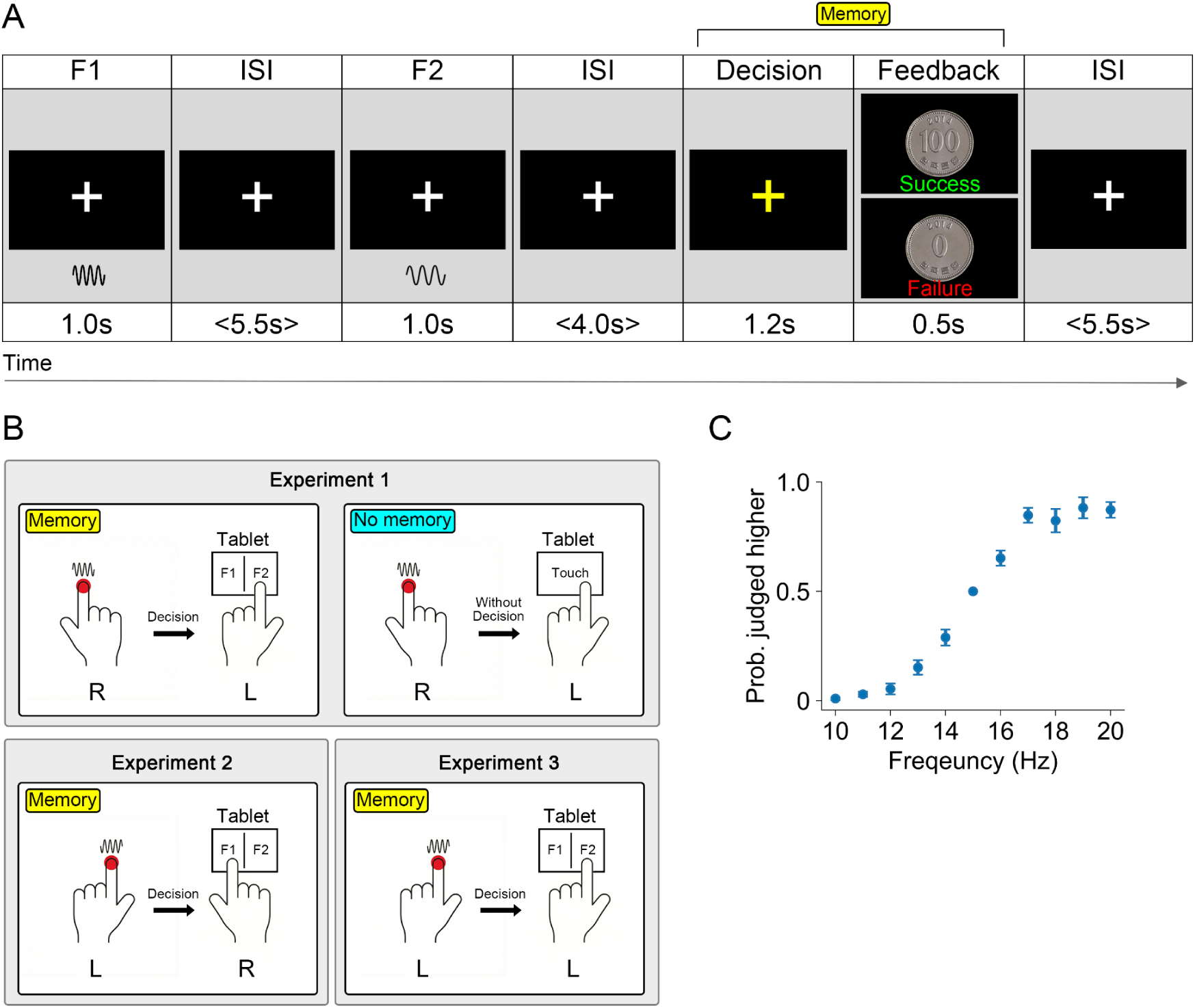
Overview and behavioral performance of the vibrotactile discrimination task. **(A)** Trial structure of the memory condition. Two sequential vibrotactile stimuli (1 s each; one fixed at 15 Hz, the other varying between 10–20 Hz) were delivered to the index finger, separated by a jittered interstimulus interval (ISI; mean 5.5 s). After a second jittered delay (mean 4.0 s), participants judged which stimulus had the higher frequency within 1.2 s of a decision cue by touching the left or right side of a tablet screen (left = first stimulation, right = second stimulation). Correct and incorrect responses were followed by images of a 100-won or 0-won coin, respectively, and each trial ended with a jittered intertrial interval (mean 5.5 s). **(B)** Experimental design across the three experiments. Experiment 1 included memory and no-memory conditions. In the memory condition, stimulation was applied to the right finger and participants indicated their decisions by touching the tablet with the left hand. In the no-memory condition, participants received identical stimulation but simply tapped the tablet with the left hand at the decision cue without comparing the stimuli. Experiment 2 tested the memory condition, where stimulation was applied to the left finger and participants indicated their decisions by touching the tablet with the right hand. Experiment 3 tested the memory condition, where stimulation was applied to the left finger and participants indicated their decisions by touching the tablet with the left hand. **(C)** Behavioral performance for the memory condition in Experiment 1. Psychometric curve showing the probability of judging the variable stimulus as higher frequency as a function of frequency (Hz). Error bars represent ± SEM across participants.

### Contralateral activation and ipsilateral deactivation during tactile stimulation

To investigate how cortical activity changes across successive tactile stimulations under the memory condition (Fig. 1A), we conducted whole-brain general linear model (GLM) analyses for stimulation of either the right (Experiment 1) or left finger (Experiment 2 and 3) (Fig. 2). In Experiment 1 (right finger stimulation), the GLM analysis for the first stimulation revealed significant activation in the contralateral primary (S1; *t*(15) = 8.47, *P* < 0.001) and secondary (S2; *t*(15) = 8.56, *P* < 0.001) somatosensory cortices, alongside deactivation in the ipsilateral S1 (*t*(15) = -9.28, *P* < 0.001). During the second stimulation, the activation pattern expanded beyond the primary sensory areas, with robust responses emerging in the bilateral S2, insular cortex, superior temporal cortex, and superior frontal gyrus (Supplementary Tables S1, S3). Notably, Experiment 2 (left finger stimulation) showed a mirrored pattern of activation and deactivation across both the first and second stimulations in the corresponding hemispheres.

**Fig. 2.**
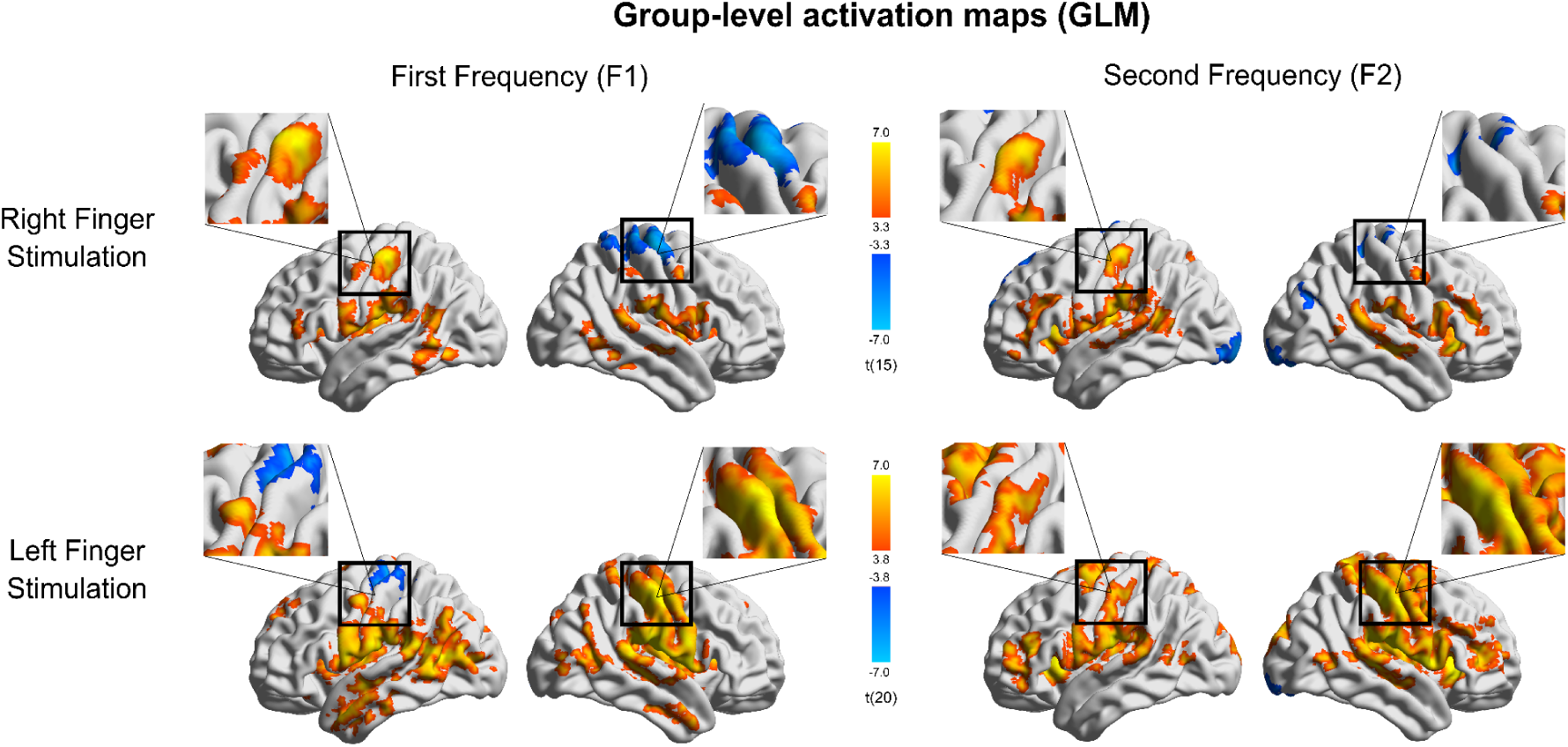
Whole-brain cortical activation during the tactile memory task. General linear model (GLM) maps illustrating BOLD responses evoked by the first and second stimulations in the memory condition for right and left finger stimulation. (uncorrected voxel-wise *P* < 0.005 for Experiment 1 and *P* < 0.001 for Experiment 2, cluster size > 40 voxels). Orange and blue colors indicate activation and deactivation, respectively.

We hypothesized that if this expanded network reflects perceptual decision-making, higher-order hubs such as the supplementary motor area (SMA) and bilateral anterior insular cortex would show selectively enhanced activation when participants compared the current stimulus with a remembered one (*11*, *16*, *17*). To dissociate this active cognitive process from sensory processing associated with stimulus presentation, we included a “no-memory” baseline condition in which participants received identical vibrotactile stimuli but were not required to maintain the stimuli in memory to compare their frequencies. Specifically, we expected these decision-making hubs to show enhanced activation during the second stimulation f2), exclusively under the memory condition. To test this, we examined the interaction between mnemonic demands, Memory (*M*) vs. No-memory (*NM*) and stimulation phase (f1 vs f2) [(M_f1_ - M_f2_) - (NM_f1_ - NM_f2_)]. Consistent with our hypothesis, the whole-brain interaction analysis revealed significant clusters prominently located within these specific decision-making hubs, SMA (peak MNI: *x* = 3, *y* = 21, *z* = 49) and bilateral anterior insula (right peak MNI: *x* = 31, *y* = 25, *z* = -5; left peak MNI: *x* = -31 *y* = 25, *z* = -3) (Fig. 3A). To further characterize this interaction while avoiding circularity, we performed post hoc analyses using independently defined atlas-based ROIs (*18*). Parameter estimates extracted from the SMA and anterior insula showed that the interaction was driven by enhanced activation during the decision phase of the memory condition (SMA: *F*(1, 15) = 22.10, *P* < 0.001; anterior insula: *F*(1, 15) = 15.15, *P* = 0.001), which sharply contrasted with the response suppression observed in the no-memory baseline (Fig. 3B). This finding suggests that the temporal transition in neural representations is not merely a consequence of successive stimulus presentation, but is dynamically modulated by the task demands of making a working memory-guided decision.

**Fig. 3.**
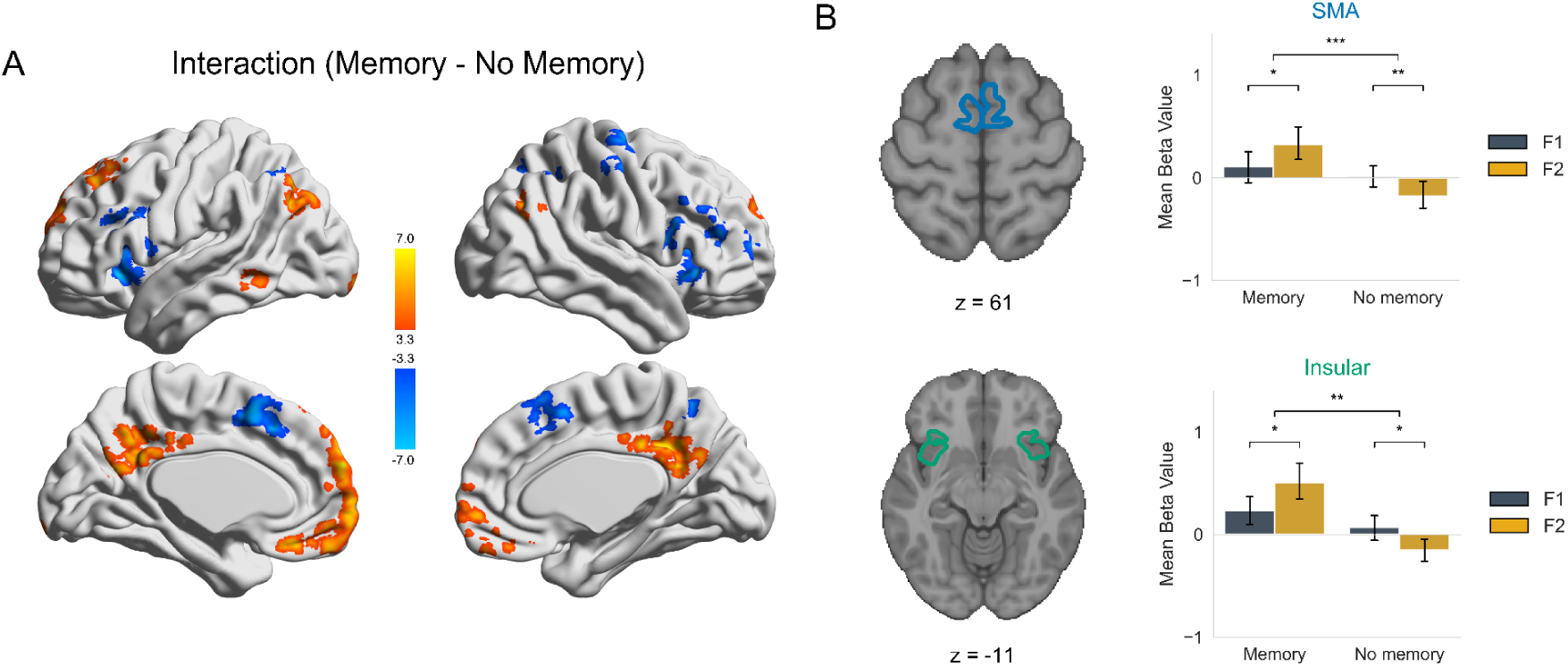
Memory demands modulate phase-dependent activation in decision-related cortices. **(A)** Whole-brain statistical map revealing regions with a significant interaction between memory context (Memory [*M*] vs. No-memory [*NM*] in Experiment 1) and stimulation phase (f1 vs f2). The map highlights areas where the temporal evolution of the BOLD response from the first to the second tactile stimulation is dependent on the memory condition, corresponding to the contrast [(M_f1_ - M_f2_) - (NM_f1_ - NM_f2_)]. Maps are thresholded at an uncorrected voxel-wise *P* < 0.005 (cluster size > 30 voxels for spatial visualization). **(B)** ROI analysis of the interaction effect. Parameter estimates (arbitrary units) were extracted from the supplementary motor area (SMA) and anterior insular cortex, defined via the HCP Glasser parcellation (MNI space). Bar plots illustrate the mean interaction effect, demonstrating enhanced frequency discrimination under the memory condition within these decision-making hubs. Error bars represent ± SEM.

### Ipsilateral activation is negatively modulated by stimulation frequency

We next investigated how tactile frequency was parametrically encoded in the sensorimotor regions using a linear mixed-effects (LME) model analysis for the whole brain (Fig. 4). For the first stimulation, frequency-dependent activity was modest and did not yield reliable clusters in the corrected whole-brain analysis (Supplementary Table S1). Nonetheless, a subtle, spatially localized modulation was observed in the ipsilateral S1 at an uncorrected threshold (p < 0.005). For visualization, we extracted BOLD responses from a spherical ROI defined around the center of mass of the ipsilateral S1 cluster. This illustrative analysis (Fig. 4, inset) showed that BOLD responses decreased with increasing stimulation frequency (*β* = -0.08, *z* = -4.33, *P* < 10^-3^), indicating frequency-dependent attenuation. During the second stimulation, this frequency-dependent suppression within the ipsilateral S1 persisted (*β* = -0.15, *z* = -5.67, *P* < 10^-5^), indicating robust and consistent parametric encoding across successive stimulation phases. Furthermore, the whole-brain LME analysis for this second stimulation revealed that this negative frequency-dependent modulation extended across a broad sensorimotor, frontoparietal, and cerebellar network. Beyond the ipsilateral S1, these regions encompassed the bilateral superior parietal lobules (SPL), dorsolateral prefrontal cortex (DLPFC), ventral premotor cortex (PMv), and cerebellar cortices (Supplementary Table S1).

**Fig. 4.**
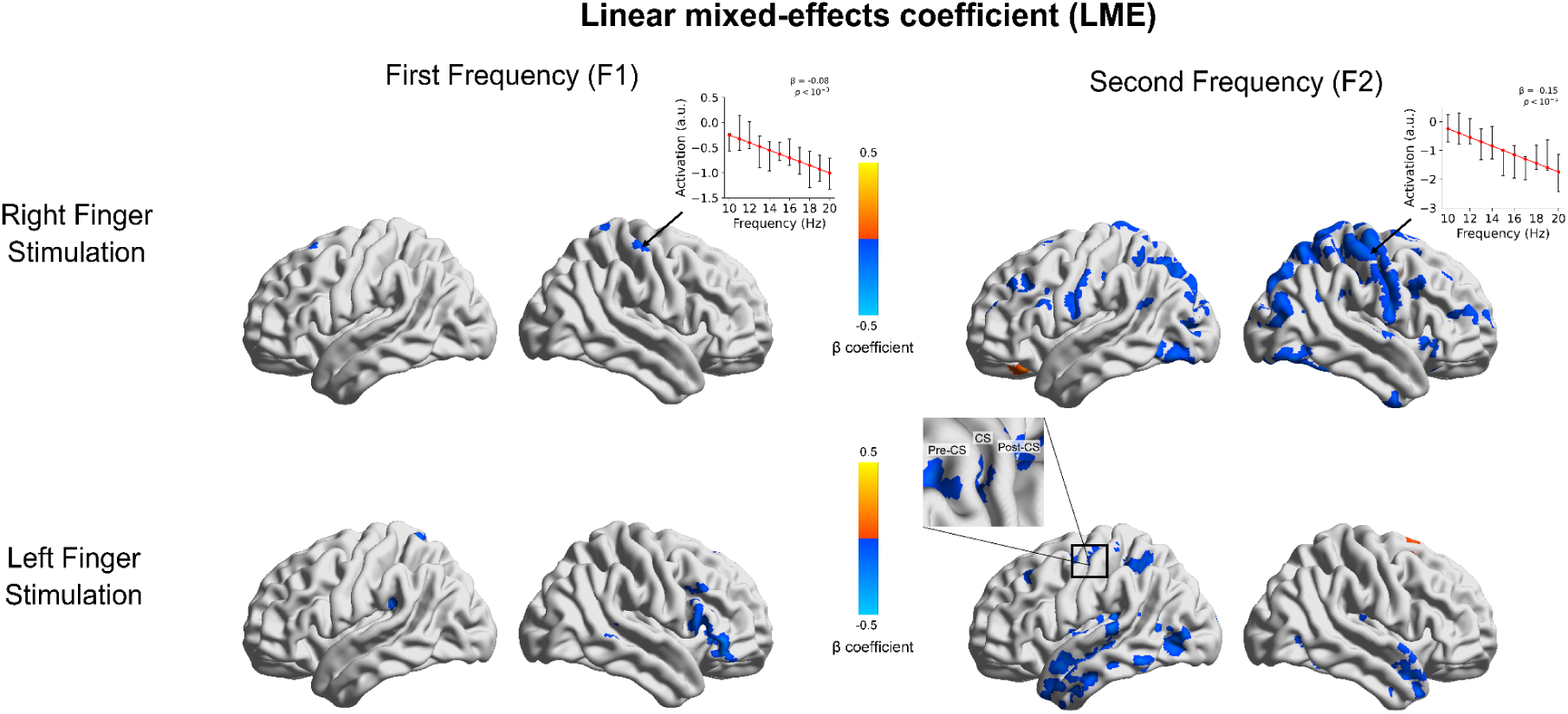
Frequency-dependent modulation of BOLD responses. Whole-brain linear mixed-effects (LME) maps demonstrate the modulation of BOLD activity across stimulation frequencies during right finger stimulation in Experiment 1 and left finger stimulation in Experiment 2 under the memory condition (uncorrected voxel-wise *P* < 0.005, cluster size > 30 voxels for spatial visualization; see Table S1,S3 for FWE-corrected clusters). Inset line plots illustrate ROI-based summaries from the ipsilateral primary somatosensory cortex (S1), displaying the mean predicted BOLD responses (± SEM) as a function of frequency. The S1 ROIs were defined using a 3-mm spherical mask centered at the center of mass of the respective significant clusters.

These findings were independently replicated in Experiment 2, in which the stimulation was delivered to the left finger. For the first stimulation, we did not observe the ipsilateral S1 modulation; instead, significant clusters emerged in the right inferior frontal cortex, right supplementary motor area (SMA), and left cerebellar cortex. During the second stimulation, the whole-brain LME analysis yielded significant clusters across a distributed network, encompassing the insula, intraparietal sulcus, temporal cortices, and subcortical structures such as the putamen and thalamus (Supplementary Table S3). Although the whole-brain LME results for Experiment 2 were less extensive, likely due to limited statistical power, and did not survive stringent cluster-level correction in the ipsilateral S1, the spatial distribution observed at a liberal threshold (*p* < 0.005, uncorrected) exhibited a spatial topography consistent with the broad network identified in Experiment 1 (Fig. 4). This strong spatial consistency, regardless of the stimulated hand, corroborates that the frequency-dependent modulation is a reproducible, system-wide phenomenon.

Together, these LME results suggest that ipsilateral S1 deactivation can vary parametrically with tactile input features, particularly during the second stimulation. This pattern is consistent with prior evidence that ipsilateral S1 negative BOLD responses during unilateral somatosensory stimulation scale with stimulus intensity and behavioral sensory suppression (*19*, *20*).

### Frequency-dependent activation does not translate into discriminable representation

While the univariate LME model analysis revealed macroscopic frequency-dependent activation, we further investigated whether tactile frequency was also represented in fine-grained multivoxel activity patterns. Using whole-brain searchlight analyses, we first examined representational dissimilarity among tactile frequencies separately within the first and second stimulations. In contrast to the univariate LME results, multivoxel pattern analyses provided little evidence for a reliable frequency representation. Specifically, whole-brain analysis revealed no significant pattern dissimilarity among stimulation frequencies within the first or the second stimulation (Supplementary Fig. S4). These findings indicate that the stimulation frequency robustly scales the overall BOLD amplitude, but does not produce discriminable activity patterns.

### Tactile discrimination is primarily represented in the ipsilateral S1

Having found no reliable frequency-specific representation within either stimulation phase, we next examined whether the representation dissimilarity was instead driven by the distinction between the first and second stimulations. For this analysis, we computed the mean dissimilarity between activity patterns evoked by the first and the second stimuli across frequency pairs actually experienced within trials (Supplementary Fig. S3). We found robust representational dissimilarity between two consecutive stimuli within the sensorimotor cortex (voxel-wise *P* < 0.001) (Fig. 5A). While this cross-phase dissimilarity was evident in both hemispheres, it was strikingly more prominent in the ipsilateral hemisphere. Specifically, the ipsilateral primary somatosensory cortex (S1) and superior parietal lobule (SPL), extending along the postcentral sulcus (poCS) and into adjacent premotor regions, exhibited markedly greater representational dissimilarity than their contralateral counterparts.

**Fig. 5.**
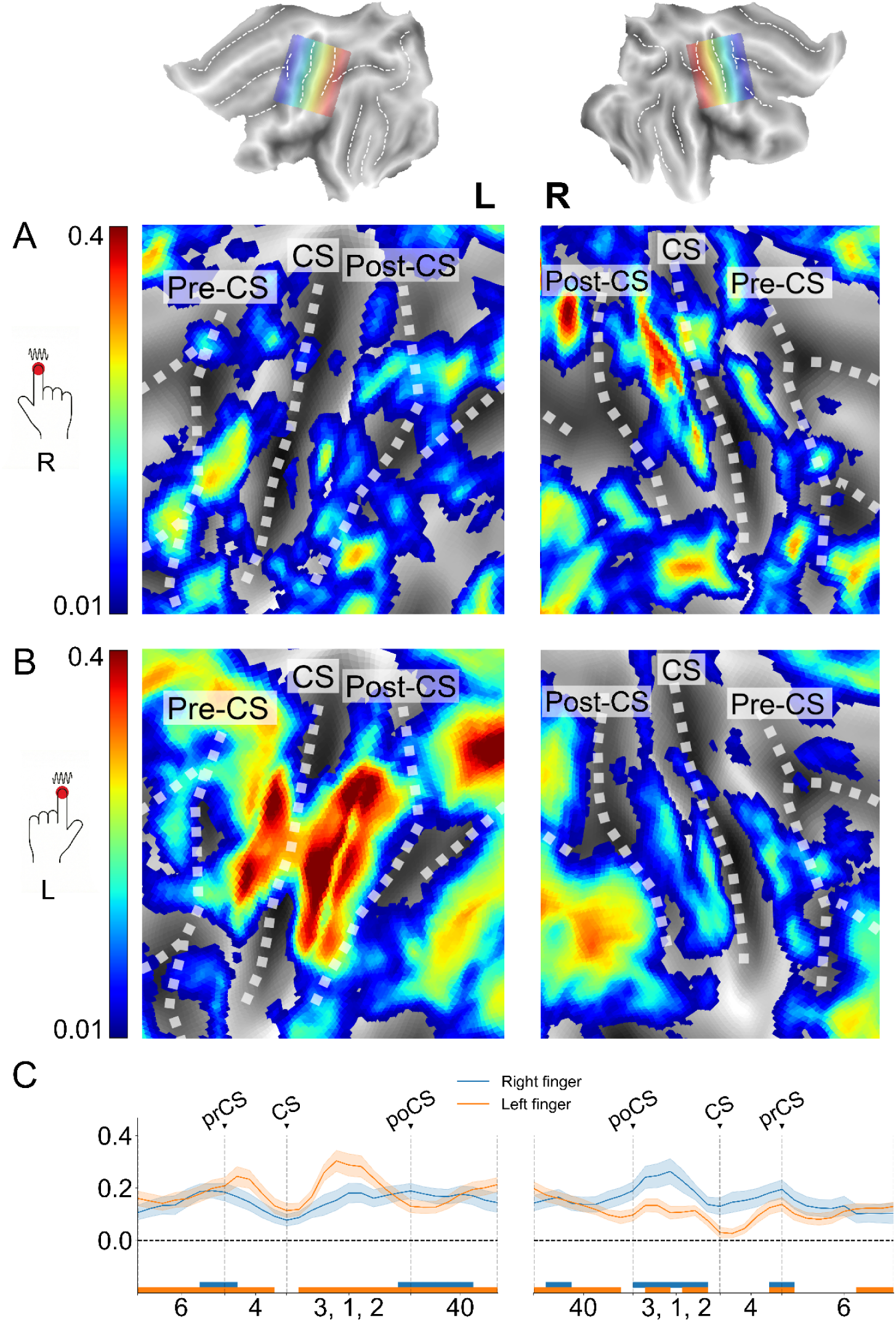
Representational dissimilarity between successive tactile stimuli (multivariate analysis). **(A)** Right-finger stimulation. Group-averaged maps of mean crossnobis dissimilarity (voxel-wise uncorrected *P* < 0.001). **(B)** Left-finger stimulation. Same as (A), but for left-finger stimulation. Note the robust dissimilarity predominantly in the ipsilateral (left) hemisphere (voxel-wise uncorrected *P* < 0.001), mirroring the pattern observed in (A). **(C)** Cortical profile plots along the sensorimotor strip. Mean representational dissimilarity is plotted from the precentral (prCS) to the postcentral sulcus (poCS) for right-finger (blue) and left-finger (orange) stimulation (group mean ± SEM). The profiles show hand-specific peaks at the poCS within each hemisphere. Horizontal bars indicate significant segments (*p* < 10⁻⁴).

We next examined whether the ipsilateral representation was also observed during left-finger stimulation. Group-averaged maps revealed a robust representational dissimilarity between the first and second stimulus in the sensorimotor regions ipsilateral to the stimulated finger (i.e., left) (voxel-wise *P* < 0.001; Fig. 5B). The representation dissimilarity was most pronounced along the poCS and extended into adjacent parietal regions, demonstrating that left-finger stimulation produced an ipsilateral activity pattern mirroring that observed during right-finger stimulation. To further delineate the spatial distribution of the activity pattern dissimilarity, we examined the representational profiles across the sensorimotor regions (Fig. 5C). These profiles revealed a robust ipsilateral bias: each stimulated hand showed greater dissimilarity in the corresponding ipsilateral hemisphere. Despite this hand-specific lateralization,, the representational profiles remained mirror-symmetric across hemispheres, with dissimilarity for both hands peaking consistently between the CS and poCS (group mean ± SEM, *P* < 10⁻⁴).

These findings suggest that distributed multivoxel activity patterns across the sensorimotor cortex do not merely encode absolute tactile frequencies but instead reflect the task-relevant distinction between two successive tactile stimuli required for perceptual decision-making.

### Representation in the ipsilateral S1 depends on cognitive context

We next examined whether this bilateral, ipsilaterally dominant dissimilarity between two stimuli was specific to memory-guided tactile decision-making or also emerged during passive stimulation without mnemonic demands. To this end, we analyzed the no-memory condition in Experiment 1, in which participants passively received the successive stimuli without subsequent decision-making. Group-averaged maps and cross-sectional cortical profiles showed that the representational dissimilarity between two stimuli was markedly attenuated in the no-memory condition compared to the memory condition across the entire ROIs (*t*(15) = 3.58, *P* = 0.002) although the dissimilarity remains significantly larger than zero (*t*(15) = 4.17, *P* < 0.001) (Supplementary Fig. S5). The reduction was particularly pronounced within the ipsilateral somatosensory and higher-order parietal regions, suggesting that the differentiation between successive tactile stimuli depends on the cognitive context in which they are processed. Thus, the ipsilateral representations appear to reflect active maintenance and comparison of tactile information, rather than an elementary sensory response to repeated stimulation.

### Ipsilateral S1 representations are not explained by motor preparation

In our primary task, participants responded using the hand contralateral to the stimulated finger. Although we minimized motor confounds using jittered inter-stimulus intervals, the representational dissimilarity in the ipsilateral cortex could, in principle, reflect preparatory signals for the responding hand. To address this possibility, we conducted a control experiment (Experiment 3) under the memory condition, in which participants received stimulation and reported their decisions with the same hand (i.e., left-finger stimulation and left-hand response).

Multivariate analyses revealed that the representational differentiation between successive stimulation phases (f1 vs f2) was preserved across the bilateral sensorimotor network. Critically, the ipsilateral S1 continued to show a robust differentiation between the first and second stimuli even when that hemisphere was not involved in preparing the motor response (peak *t*(15) = 6.04, *P* < 10⁻⁴) (Supplementary Fig. S6). These findings make it unlikely that the ipsilateral f1 - f2 representation can be explained by motor preparation alone. Rather, they further strengthened our interpretation that the ipsilateral sensorimotor regions contribute to the task-relevant differentiation of successive tactile stimuli, which is required for perceptual decision-making.

## DISCUSSION

This study investigates how the human brain represents perceived vibrotactile differences within somatosensory networks. Using univariate and multivariate fMRI analyses, we show that the ipsilateral primary somatosensory cortex (S1), compared to the contralateral counterpart, primarily contains task-relevant information when comparing two consecutive vibrotactile stimuli. Our findings challenge the traditional view that tactile perception is supported predominantly by contralateral somatosensory information processing.

At the univariate level, we first confirmed the classical contralateral dominance of responses to unilateral tactile stimulation. Both the first and second stimuli evoked robust activation in contralateral S1 and concurrent deactivation in ipsilateral S1. This is consistent with previous findings of interhemispheric inhibition during unilateral input (*1*, *2*, *4*). Crucially, our linear mixed-effects (LME) model analysis revealed a significant negative linear effect of frequency during the second stimulus period in the ipsilateral S1. This macroscopic amplitude modulation is consistent with previous fMRI findings showing frequency-dependent decreases in BOLD activity in both contralateral and ipsilateral S1 during vibrotactile stimulation (*21*). This frequency-dependent suppression may reflect transcallosal inhibition from contralateral S1. By reducing activity in the unstimulated hemisphere, this mechanism could help maintain sensory contrast (*13*, *20*). Its stronger expression during the second stimulation may further suggest task-dependent modulation during memory-guided comparison. In contrast, our RSA analysis showed little evidence of stimulation frequency representation at the multivariate pattern level (*22*). One possible explanation is that our task used a relatively narrow frequency range, limited to 10-20 Hz, which falls within the flutter domain and therefore evokes relatively similar perceptual and neural responses. In contrast, previous work tested a much broader range (20–200 Hz), spanning distinct perceptual domains including flutter and vibration, which are associated with distinct underlying neural pathways (*22*).

The multivariate results provided little evidence for separable representations of individual stimulation frequencies. Instead, activity patterns more strongly differentiated the first and second stimuli during tactile decision-making. Importantly, this representational dissimilarity was greater in the ipsilateral sensorimotor regions than in the corresponding contralateral regions.

Taken together, our results from the univariate GLM and multivariate RSA analyses point to complementary aspects of ipsilateral S1 processing. The GLM analysis showed that tactile frequency was represented in local response amplitude, indicating frequency-related coding at the voxel-wise level. In contrast, the RSA analysis showed that multivoxel activity patterns provided little evidence for stimulus frequency, but instead revealed robust pattern differentiation between the two stimuli during tactile decision-making. These findings are consistent with previous evidence from animal studies demonstrating a shift from focal activation during the initial stimulus to a more distributed representation during the second stimulus (*10*, *12*, *23*). This progression suggests a transition from elementary sensory processing to higher-order representations that facilitate subsequent comparison and decision-making (*24*). Thus, ipsilateral S1 BOLD responses may represent tactile information at two levels: local amplitude modulation related to stimulus frequency and distributed patterns related to the comparison of current input with a remembered sensory trace (*10*, *25*).

The importance of this comparative computation is further supported by the results from the no-memory control condition. In this condition, participants received two successive tactile stimuli without the requirement to maintain or compare them. This allowed us to dissociate active cognitive processing from sensory responses to stimuli. Our findings suggest that mnemonic demands modulated both univariate responses and multivariate activity patterns. When working memory was not required, the targeted f2-related enhancement in the SMA and bilateral anterior insula was absent. Furthermore, at the univariate level, the ipsilateral suppression in S1 was markedly attenuated and did not survive cluster-level correction (Supplementary Fig. S2 and Supplementary Tables S2). Concurrently, at the multivariate level, the representational dissimilarity (f1 vs f2) within the ipsilateral somatosensory and higher-order parietal regions was substantially reduced (Supplementary Fig. S5). Together, this converging evidence suggests that the neural differentiation between successive tactile stimuli is not merely stimulus-driven response to repeated tactile inputs. Rather, it reflects information processing related to working memory-guided decision-making (*12*, *13*).

The memory-dependent modulation of ipsilateral representations and their hemispheric symmetry suggest that tactile cognition may rely on coordinated interhemispheric interactions (2, 4). This recruitment of the ipsilateral hemisphere is unlikely to be a mere by-product of unilateral sensory processing. Instead, our findings suggest that task demands refine sensory codes through active cortical interactions (10, 29). Specifically, this enhancement may be supported by top-down feedback from association cortices, such as parietal regions, to facilitate perceptual discrimination (8, 12). By conveying information about decisions and expectations to early sensory regions, these higher-order areas may bias somatosensory activity toward the features most relevant to current goals. Dynamic interhemispheric coupling may thus allow the brain to flexibly allocate representational resources based on cognitive demands (2, 30). Ultimately, these findings support a view of tactile perception as a bilateral, context-dependent process (12, 13). By showing that ipsilateral somatosensory areas modulate sensory representations when information must be compared, this evidence moves beyond the traditional model of purely contralateral coding and highlights interhemispheric coordination as an important component of tactile decision-making (24).

Importantly, the ipsilateral f1-f2 representation was not specific to one hand or one hemisphere. Regardless of stimulated fingers, right or left, a similar pattern emerged primarily in the ipsilateral sensorimotor regions. Although minor topographic variations existed, the overall spatial distribution and tuning profiles were consistent across hemispheres. This symmetry suggests that ipsilateral representational coding is not an idiosyncratic feature of one hemisphere, but rather a shared organizational feature of the bilateral somatosensory network. Furthermore, to rule out the possibility that these ipsilateral representations were confounded by motor planning or response execution, we verified the results in a control condition in which participants responded using the same hand that received the stimulation. Even under this setup, the ipsilateral S1 continued to exhibit significant phase-specific multivariate pattern differentiation (Supplementary Fig. S6). Together, these findings suggest that ipsilateral sensorimotor regions contribute to the comparison of successive tactile stimuli during tactile decision making, arguing against a purely motor explanation for the ipsilateral activity.

Several limitations should be considered when interpreting these findings. First, Experiment 2 showed weaker whole-brain effects than Experiment 1. The LME analysis revealed a less extensive network and lower statistical sensitivity, which may be partly explained by missing frequency conditions in several runs. Missing or unevenly sampled frequencies likely reduced the stability of the frequency-dependent model.

A second methodological limitation concerns the FIR modeling approach. The frequency-specific FIR analysis was restricted to delays 1–3 because of the limited number of trials per condition. This restriction appears unlikely to have strongly affected the main findings, because the observed tactile-evoked BOLD response peaked within this window and decreased thereafter. Consistent with this, a collapsed control model that estimated delays 0–4 showed a broadly similar spatial pattern when delays 1–3 were summed. Thus, the delay restriction did not appear to substantially alter the main effects.

A further limitation is that the observed ipsilateral f1-f2 dissimilarity did not reliably correlated with discrimination accuracy. A more explorative whole-brain searchlight RSA did not identify any significant regions where representational dissimilarity explained behavioral performance. Thus, the ipsilateral S1 representation should not interpreted as a direct neural readout of behavioral performance, as demonstrated in Figure 1C. Rather, it may reflect an intermediate process of comparing two successive stimuli that is subsequently intergrated with downstream decision-making processes.In conclusion, our findings challenge a well-established contralateral view of tactile processing. Although unilateral tactile stimulation produced the classical pattern of contralateral S1 activation and ipsilateral deactivation, tactile decision-making that required differentiating the first and second stimuli engaged ipsilateral S1 more strongly than contralateral S1. While the current study focused on vibrotactile frequency discrimination, it remains unclear whether similar ipsilateral representations would be observed for other somatosensory features, such as texture, spatial location, or tactile patterns (11, 26–28). Future studies using tactile pattern discrimination tasks could clarify whether ipsilateral S1 contributes broadly to tactile working memory and decision-making, or whether its role is more specific to frequency-dependent temporal processing.

## MATERIALS AND METHODS

### Participants

*Experiment 1.* Eighteen healthy young adults were recruited. All participants reported no history of neurological or psychiatric disorders and were not taking centrally acting medications. All participants were right-handed and reported normal tactile and visual sensitivity. Two participants were excluded (one from data analysis due to chance-level task performance and the other for discontinuing participation due to claustrophobia), leaving 16 participants (6 females; mean age ± SD = 22.1 ± 1.8 years) for analysis.

*Experiment 2*. A separate group of 26 healthy adults (12 females; mean age ± SD = 22.9 ± 2.5 years) participated in Experiment 2. Five participants were excluded (four due to chance-level task performance and one due to failure of medical screening), leaving 21 participants (9 females, 12 males) for analysis. The selection criteria are identical to those used in Experiment 1.

*Experiment 3*. Thirteen healthy adults (7 females; mean age ± SD = 22.6 ± 4.0 years) participated in Experiment 3. Notably, four of these participants had previously participated in Experiment 2, whereas the remaining nine were newly recruited for this experiment. All participants met the same inclusion criteria as in Experiments 1 and 2.

The study protocol in both experiments was approved by the Institutional Review Board of Sungkyunkwan University (IRB No. 2018-05-003-038) and Hanyang University (IRB No. HYUIRB-202508-002-1). All participants in both experiments provided written informed consent in accordance with the Declaration of Helsinki.

### Task procedures and experiment design

Vibrotactile stimuli (1 s duration) were delivered sequentially to either the right (Experiment 1) or left (Experiment 2) index finger using an MR-compatible stimulator (mini PTS-T1, Dancer Design, UK). On each trial, one stimulus had a fixed frequency of 15 Hz, while the other varied between 10 and 20 Hz in 1-Hz increments (excluding 15 Hz), ensuring that the two frequencies always differed. Stimuli were separated by a jittered interstimulus interval between 3.5 s and 7.5 s (mean = 5.5 s), followed by another jittered delay between 2.0 s and 6.0 s (mean = 4.0 s), followed by a 1.2 s decision period.

In Experiment 1, vibrotactile stimuli were applied to the right index finger, and participants performed two task conditions: “memory” and “no-memory” conditions. In the “memory” condition, participants judged which of the two stimuli had the higher frequency.

They responded by touching the left or right corner of the tablet screen to indicate whether the first or second stimulus, respectively, had the higher frequency with their left hand within 1.2 s. Correct and incorrect responses were followed by images of a 100-won or 0-won coin, respectively, and participants were informed that monetary compensation would be proportional to their accumulated reward (*17*, *26*, *27*). In the no-memory condition, participants received the same tactile stimuli but were instructed to tap the tablet during the decision period without comparing the stimuli or judging their frequencies. Feedback images were presented at random, regardless of the participants’ responses. Each condition consisted of six runs of 20 trials each, with pseudorandomized frequency pairs. To minimize carryover of comparison strategies, all no-memory sessions were completed first, followed by acquisition of the T1-weighted anatomical scan and the memory sessions.

In Experiment 2, vibrotactile stimuli were applied to the left index finger under the memory condition. Participants responded by touching a tablet screen with their right hand, while the task structure and feedback procedures were identical to those in Experiment 1. Experiment 2 consisted of three runs with 40, 30, and 30 trials, respectively. Due to differences in the run structure and available frequency combinations, some frequency pairs present in Experiment 1 were not included in Experiment 2 (run 2: first stimulus at 19 Hz and first stimulus at 20 Hz; run 3: first stimulus at 14 Hz, first stimulus at 18 Hz, and second stimulus at 19 Hz). These missing pairs were excluded from subsequent representational similarity analyses.

Experiment 3 adopted the trial structure of Experiment 1, consisting of six runs of 20 trials each, while utilizing left index finger stimulation, as in Experiment 2. The key modification in Experiment 3 was that participants responded with the thumb of the stimulated left hand instead of the contralateral hand. This setup ensured that both sensory input and motor response were localized to the same limb. Except for the differences noted above, all task parameters were kept consistent across experiments. All tasks were programmed using MATLAB (Mathworks, Inc.).

### MRI data acquisition

We conducted experiments at the Center for Neuroscience Imaging Research at Sungkyunkwan University, using a 3T MRI scanner equipped with a 64-channel head coil. Earplugs and form pads were employed to reduce noise and minimize head motion during scans. For all the functional scans, we used echo planar imaging (EPI) sequence with parameters set as follows: repetition time (TR) = 2000 ms; voxel size = 2.0 x 2.0 x 2.0 mm; echo time (TE) = 30 ms; field of view (FOV) = 200 x 200 mm; flip angle (FA) = 90°; slice thickness = 2.0 mm; matrix size = 100 x 100 x 90 voxels; EPI images were captured in the anterior-to-posterior direction. For Experiment 1, twelve functional runs were acquired, comprising six for a no-memory task and six for a memory task. For Experiment 2, three functional runs were acquired exclusively for a memory task. For Experiment 3, six functional runs were acquired exclusively for a memory task. Additionally, a whole-brain T1-weighted anatomical scan was performed using a magnetization-prepared rapid acquisition with gradient echo (MPRAGE) sequence with the following settings: repetition time (TR) = 2300 ms; voxel size = 1.0 x 1.0 x 1.0 mm; echo time (TE) = 2.28 ms; field of view (FOV) = 192 x 256 mm; flip angle (FA) = 8°; slice thickness = 1.0 mm; matrix size = 192 x 256 x 256 voxels.

### Analysis

#### Behavioral data analysis

Each trial was scored as “comparison frequency judged higher” or “not higher” relative to the 15 Hz reference. For each of participants and comparison frequencies (10–20 Hz), we computed the proportion of “judged higher” responses and then averaged across participants to obtain group-level discrimination accuracy. We quantified tactile discrimination performance by fitting a psychometric function to these proportions with a cumulative Gaussian function. Then, we characterized discrimination sensitivity by the slope of the fitted curve, which provided a quantitative measure of perceptual sensitivity, and estimated the just noticeable difference (JND) as the frequency difference corresponding to a 75% response probability (*15*). We calculated the standard error of the mean (SEM) across participants and displayed it as error bars in Fig. 1C. The figure also shows the fitted psychometric curve, illustrating how the probability of judging the comparison frequency as higher increased with frequency.

#### Preprocessing of fMRI data

Anatomical and functional imaging data were preprocessed following a standard procedure guided by AFNI’s afni_proc.py (*28*, *29*). Initially, DICOM files were converted to NIfTI forma t(*30*). T1-weighted and EPI images were processed to ensure proper file conversion. The preprocessing pipeline included several key steps. Transient signal spikes in the time series were attenuated using the (3dDespike) function. Slice-timing correction was performed with (3dTshift) using quintic interpolation, and susceptibility-related EPI distortions were corrected using forward and reverse phase-encoded EPI images. Motion correction was applied using (3dvolreg). Functional images were aligned to the anatomical T1 volume with align_epi_anat.py, and non-linear transformation to MNI space was achieved using auto_warp.py. An EPI volume extent mask was applied to exclude invalid voxels. Spatial smoothing was conducted using a Gaussian kernel with a 4-mm full-width at half-maximum using the (3dmerge) function. The time series were scaled to a mean of 100 using the (3dTstat) and (3dcalc) functions. Regression analysis, performed with (3dDeconvolve), included motion parameters and their temporal derivatives as nuisance regressors, and volumes with excessive motion or outlier fractions were censored. Residual time series were then estimated using (3dTproject). This comprehensive preprocessing pipeline ensured the fMRI data were prepared accurately and efficiently for subsequent analysis, enabling robust and reliable neuroimaging results.

#### Whole-brain GLM analysis

We conducted a whole-brain general linear model (GLM) analysis to identify regions related to vibrotactile stimulation (*31*). For the subject-level analysis, we modeled the hemodynamic response using a finite impulse response (FIR) basis with delays of 1–3 TRs (*32*). This temporal window was chosen to capture the expected BOLD response to tactile stimulation while reducing potential temporal overlap with responses to subsequent events. Because each specific frequency pair occurred exactly once within a run, including a wider finite impulse response window for individual frequency regressors could have reduced design efficiency and produced unstable beta estimates due to increased regressor collinearity. Beta estimates from delays 1–3 were summed to obtain a single parameter estimate per event. The design matrix additionally included six motion parameters, polynomial drift regressors (up to fourth order), and event regressors for decision and feedback onsets.

Two models were examined. In the first model, beta weights were estimated separately for the first and second stimulation events, with frequency collapsed, to identify regions sensitive to stimulation phase. In Model 2, beta weights were estimated separately for each comparison frequency from 10 to 20 Hz in 1-Hz steps and for each stimulation phase, yielding 22 frequency-by-phase regressors. This model allowed us to examine frequency-dependent responses as a function of stimulation phase.

Subject-level beta maps were averaged across runs and submitted to random-effects group analyses. Statistical significance was assessed using two-sided voxelwise one-sample *t*-tests (*28*). Correction for multiple comparisons was performed using cluster-level thresholding based on Monte Carlo simulations of spatial noise smoothness within the whole-brain gray-matter mask (26-neighbor connectivity criterion; FWE correction) (*33*, *34*). For visualization only, the resulting t-statistic maps were displayed using experiment-specific uncorrected voxelwise thresholds: P < 0.005 for right-finger stimulation in Experiment 1 and P < 0.001 for left-finger stimulation in Experiment 2, with a cluster-extent threshold of 40 voxels (Fig. 2). For standardized reporting, the corresponding supplementary tables (Supplementary Tables S1–S3) were reported using a uniform voxelwise cluster-forming threshold of P < 0.001, and peak t-values from clusters surviving cluster-level FWE correction were converted to equivalent Z-scores.

#### Whole-brain interaction analysis

To investigate the interaction between cognitive demands and stimulation phase, we performed an additional whole-brain analysis using AFNI’s 3dMVM (Multivariate Modeling) (*35*). We tested the interaction effect between memory context (*M* vs. *NM*) and stimulation phase (f1 vs f2) using the specific contrast [(M_f1_ - M_f2_) - (NM_f1_ - NM_f2_)]. Multiple comparison correction for the interaction map followed the identical Monte Carlo simulation parameters within the whole-brain mask, and for spatial visualization, the interaction *t*-statistic map was thresholded at an uncorrected voxel-wise *P* < 0.005 with a minimum cluster extent of 30 voxels.

#### Linear mixed-effects analysis

To assess frequency-dependent modulation of BOLD activity across participants, we performed voxelwise linear mixed-effects (LME) modeling using AFNI’s 3dLMEr (*36*). Beta estimates from the subject-level GLM (Model 2: frequency-by-phase conditions, 10–20 Hz in 1-Hz steps) were entered into the model. The fixed effect of interest was stimulus frequency (Freq), with random intercepts for participants (Subj) and nested random effects for runs within participants (Subj:Run). The model was formulated as:

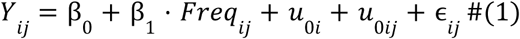

where *Y*_*ij*_ denotes the BOLD response for participant *i* in run *j*, β_1_ reflects the fixed frequency effect, *u*_0*i*_ and *u*_0*ij*_ represent random intercepts for participant and run nested within participant, and ɛ_*ij*_ denotes the residual error.

Statistical significance was assessed using Wald Z-statistics derived from the LME model under the asymptotic normal approximation. To correct for multiple comparisons, the spatial autocorrelation of residuals was estimated from the subject-level models. This estimation was used to determine cluster-size thresholds via Monte Carlo simulations of spatial noise smoothness within the whole-brain mask (26-neighbor connectivity criterion), ensuring family-wise error (FWE) correction. While cluster-level FWE correction was strictly applied to define statistical significance for all reported coordinates (Supplementary Table S1-S3), brain maps in the figures were thresholded at an uncorrected voxel-wise *P* < 0.005 to visualize the overarching spatial trends and extended network topographies.

#### Region-of-interest (ROI) analyses

To visualize frequency-dependent trends within the ipsilateral primary somatosensory cortex (S1), we performed an ROI analysis for visualization purposes only, without drawing independent statistical inferences from the extracted estimates. The ipsilateral S1 ROI was defined as a 3-mm-radius spherical mask centered at the center of mass of the cluster identified in the first-stimulation frequency contrast. We then applied the same LME model to beta estimates extracted from this ROI and plotted the predicted frequency–BOLD relationship based on the fixed effect of frequency. This analysis provided a visual summary of the overall frequency-dependent response pattern in ipsilateral S1.

Furthermore, to characterize the interaction effect in higher-order decision-related regions while avoiding circularity, we performed independent post-hoc ROI analyses using structurally defined atlas parcels. The SMA-related ROI was defined from the Supplementary and Cingulate Eye Field parcels of the HCP Glasser parcellation in MNI space, and the bilateral anterior insula ROI was defined by combining the Anterior Agranular Insula Complex and Anterior Ventral Insular Area parcels in both hemispheres (*18*). For each participant, these ROI masks were resampled to the corresponding beta images using nearest-neighbor interpolation, and mean beta estimates were extracted across all voxels within each ROI. These estimates were extracted separately for the four specific conditions, comprising M_f1_, M_f2_, NM_f1_ and NM_f2_. The extracted beta values were then subjected to post-hoc paired *t*-tests to visually and statistically evaluate the specific phase-dependent differences within each memory context and the overall interaction effect.

#### Representational similarity analysis

For Experiment 1, we computed local representational dissimilarity using a voxel-centered searchlight (6-mm radius; 93 voxels) implemented as spherical ROIs spanning the whole brain within the brain mask (*14*, *37*). For each spherical searchlight and participant, we built a representational dissimilarity matrix (RDM) using cross-validated Mahalanobis (crossnobis) distances (*38*) across the 22 conditions (11 comparison frequencies from 10 to 20 Hz in 1 Hz steps by two stimulus phases, first and second). To isolate the effect of cognitive context, these RDMs were constructed separately for the memory and no memory conditions. Cross-validation was performed across independent runs to ensure unbiased distance estimation (*39*) with noise normalization using residual covariance from the subject-level GLM with Ledoit–Wolf shrinkage (*40*).

From these RDMs, we computed two complementary dissimilarity metrics. First, to test whether tactile frequency was represented within a given stimulation phase (first or second), we computed within-phase frequency dissimilarity by averaging crossnobis distances between different frequency conditions separately within the first and second stimulations. This analysis tested whether multivoxel patterns reliably distinguished tactile frequencies without comparing across stimulus phases. Second, to examine representational differentiation between successive stimulations, we computed a reference-anchored cross-phase dissimilarity metric. Instead of averaging all off-diagonal entries, we computed mean dissimilarity as the mean crossnobis distance across the 20 reference-anchored pairs involving the 15 Hz reference. Specifically, we averaged: (i) pairs between 15 Hz (first) and all f Hz (second) where f ≠ 15, and (ii) pairs between all f Hz (first) and 15 Hz (second) where f ≠ 15. This metric aligns with the task’s reference-based discrimination structure and excludes non-existent pairs. The complete set of reference-anchored pairs is illustrated in Supplementary Fig. S3. Participant-level dissimilarity maps were submitted to two-sided one-sample *t* tests for a group-level analysis. For visualization purposes, the resultant group-level maps were displayed at an uncorrected voxel-wise threshold of *P* < 0.001 with a color scale capped at a dissimilarity value of 0.4 (Figs. 5A and 5B).

For Experiment 2, the same analysis pipeline was applied, but several frequency conditions used in Experiment 1 were missing in one or more runs. Consequently, reference-anchored pairs involving 14 Hz, 19 Hz, and 20 Hz (first-stimulus side) or 18 Hz and 19 Hz (second-stimulus side) could not be computed and were excluded from the RDM. Dissimilarity estimates for Experiment 2 were therefore calculated exclusively using the remaining valid reference-anchored pairs.

Furthermore, to investigate neural representations in the absence of contralateral motor demands, we applied the identical analysis pipeline used for Experiment 1 to the data from Experiment 3. In this experiment, stimulation was applied to the left finger and participants responded using the same left hand. Because the full set of frequency conditions was successfully acquired, the representational dissimilarity matrix and the reference anchored cross-phase dissimilarity metric were computed using all 20 pairs, exactly as performed in Experiment 1.

#### Cortical profile analysis

To examine spatial variations in representational dissimilarity across the sensorimotor cortices, we performed a cortical profile analysis using mean dissimilarity maps(*41*, *42*). The cortical trajectory used for profile extraction is shown in Fig. 5A and encompasses Brodmann areas 6, 4, 3/1/2, and 40, covering the precentral, central, and postcentral sulcal regions. Surface-projected (vertex-wise) values were sampled along this anatomically defined path and uniformly resampled into 30 equally spaced points. At each sampling position, values were averaged across participants to generate distinct representational profiles, with shaded areas denoting the group-level SEM. To illustrate hand-specific regional recruitment within the memory condition, values were averaged separately for right-finger and left-finger stimulations (Fig. 5C). To evaluate the effect of cognitive context, values from the right-finger stimulation were averaged separately for the memory and no-memory conditions (Supplementary Fig. S4). Finally, the corresponding cortical profile demonstrating the phase-specific representational geometry for Experiment 3 is provided in Supplementary Fig. S6. Across all profile plots, horizontal bars indicate vertex segments showing significant positive dissimilarity (*P* < 10^-4^).

## Acknowledgments

Neuroimaging was performed at the Center for Neuroscience Imaging Research located at Sungkyunkwan University, supported by the Institute for Basic Science.

## Data and code availability

All MRI data used in this study are archived in the Hanyang University Network Attached Storage (NAS). Upon reasonable request, the corresponding author can provide all the dataset and codes related to this study.

## Author contributions

S.K. conceived the idea presented in the study. S.K., H.Y., and S.P designed the experiment. H.Y. and S.P. performed the experiment. H.Y. performed formal analysis; S.Y. supported multivariate analysis and visualization of results; H.Y. and S.K. wrote the manuscript. S.K. supervised the entire procedure of the study.

