## Supplementary Materials for "Ipsilateral somatosensory cortex contains task-relevant tactile representations despite contralateral activation dominance"

Hyeree Yoon et al.

This PDF file includes:
Figs. S1 to S6
Tables S1 to S3

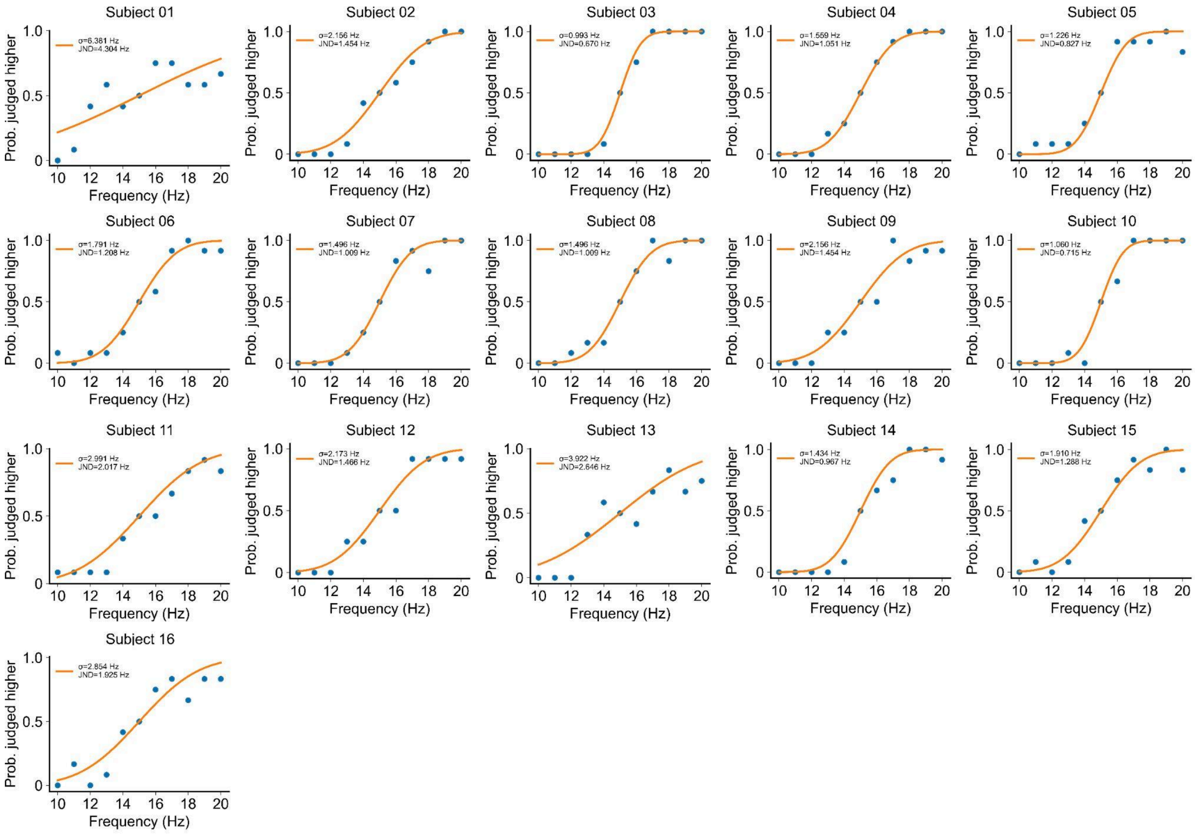


**Fig. S1. Individual behavioral performance in the vibrotactile discrimination task (Memory condition in Experiment 1).** Behavioral performance. Psychometric curves illustrate the probability of judging the variable stimulus as having a higher frequency as a function of stimulus frequency (Hz) for 16 individual participants.

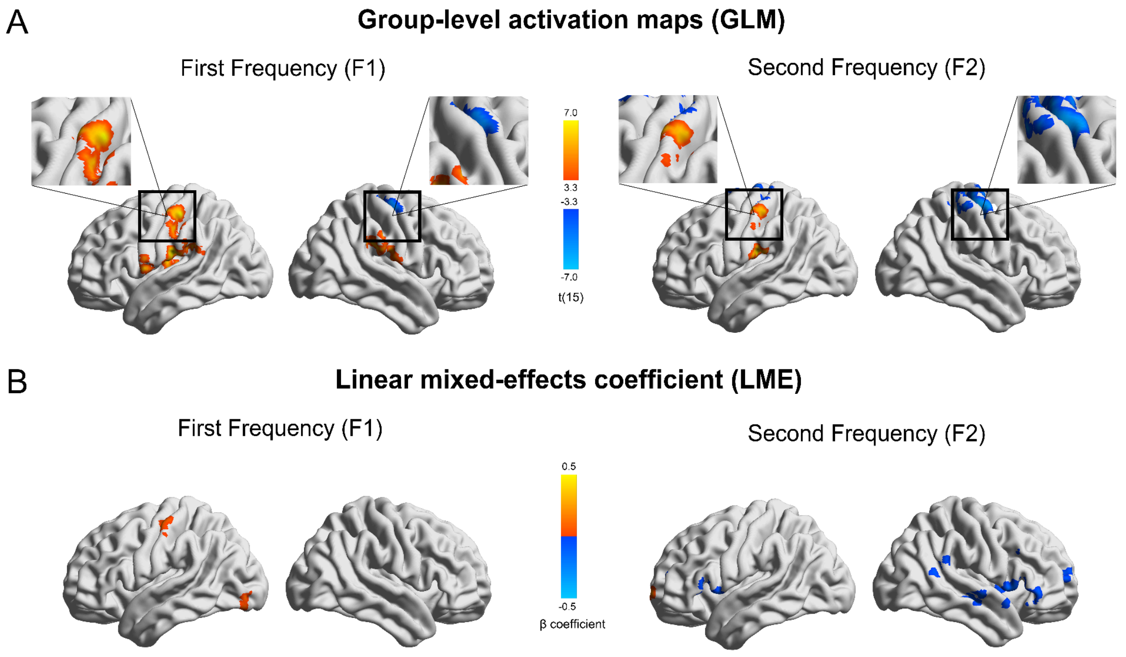


**Fig. S2. GLM and LME based analyses of frequency-dependent BOLD responses (No memory condition). (A)** Whole-brain GLM results showing cortical activation during the first and second stimulations in the no memory condition (uncorrected voxel-wise $P < 0.005$, cluster size > 40 voxels). Colors indicate activation and deactivation, respectively. **(B)** Whole-brain voxel-wise linear mixed-effects (LME) results showing frequency-dependent modulation of BOLD activity in the no memory condition (voxel-wise $P < 0.005$, cluster size > 40 voxels).

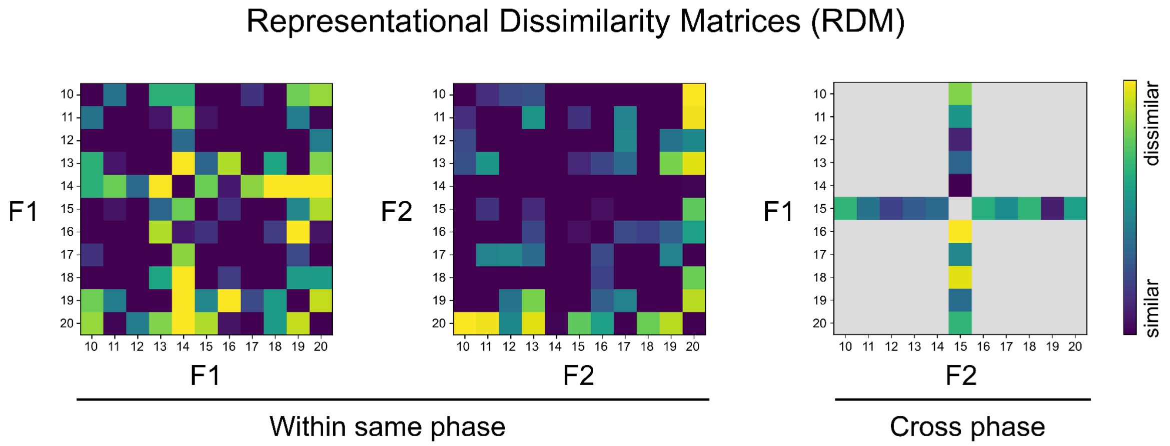


**Fig. S3. Representational Dissimilarity Matrices.** These matrices are sample images provided for illustrative purposes. The left matrix shows within phase comparisons for the first stimulation F1. The middle matrix shows within phase comparisons for the second stimulation F2. The right matrix shows cross phase comparisons between F1 and F2. For the cross phase analysis only the 20 pairs including the 15 Hz reference frequency are used. Grey areas indicate excluded pairs. The color scale shows the degree of dissimilarity.

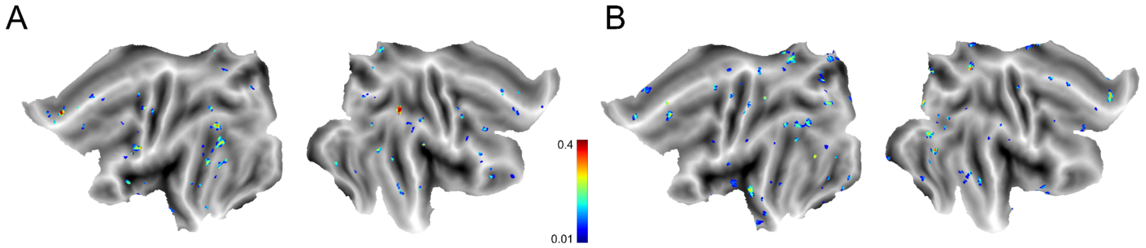


**Fig. S4. Whole brain representational dissimilarity within stimulation phases. (A)** Group averaged map of mean crossnobis dissimilarity within the first stimulation (voxel wise uncorrected *P* < 0.01). **(B)** Group averaged map of mean crossnobis dissimilarity within the second stimulation (voxel wise uncorrected *P* < 0.01).

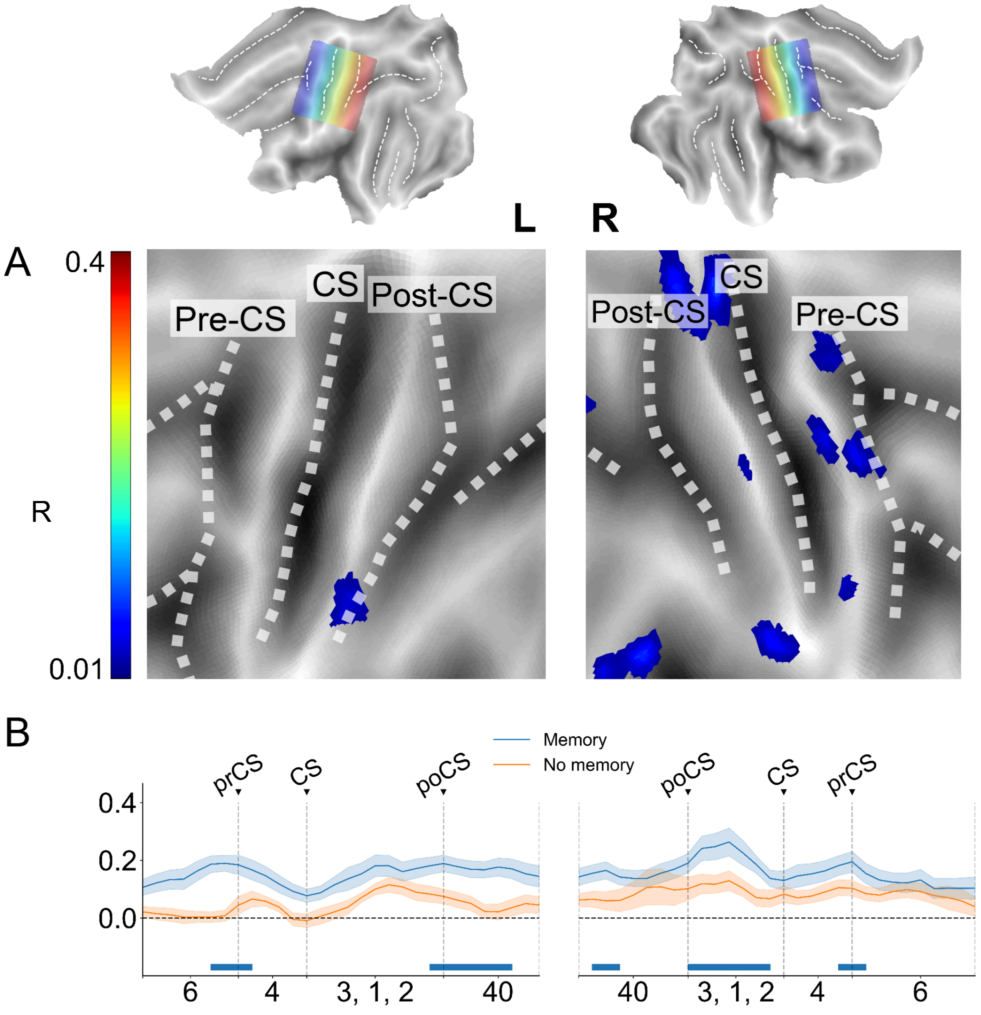


**Fig. S5. Representational dissimilarity across hemispheres for right finger stimulation in no memory condition. (A)** Group averaged map of mean crossnobis dissimilarity for right finger stimulation in the no-memory condition (voxel wise uncorrected *P* < 0.001). **(B)** Cortical profile plots along the sensorimotor strip. Mean representational dissimilarity is plotted from the precentral (prCS) to the postcentral sulcus (poCS) for right finger stimulation in the memory (blue) and no-memory (orange) conditions (group mean ± SEM). Horizontal bars indicate significant profile segments ($p < 10^{-4}$).

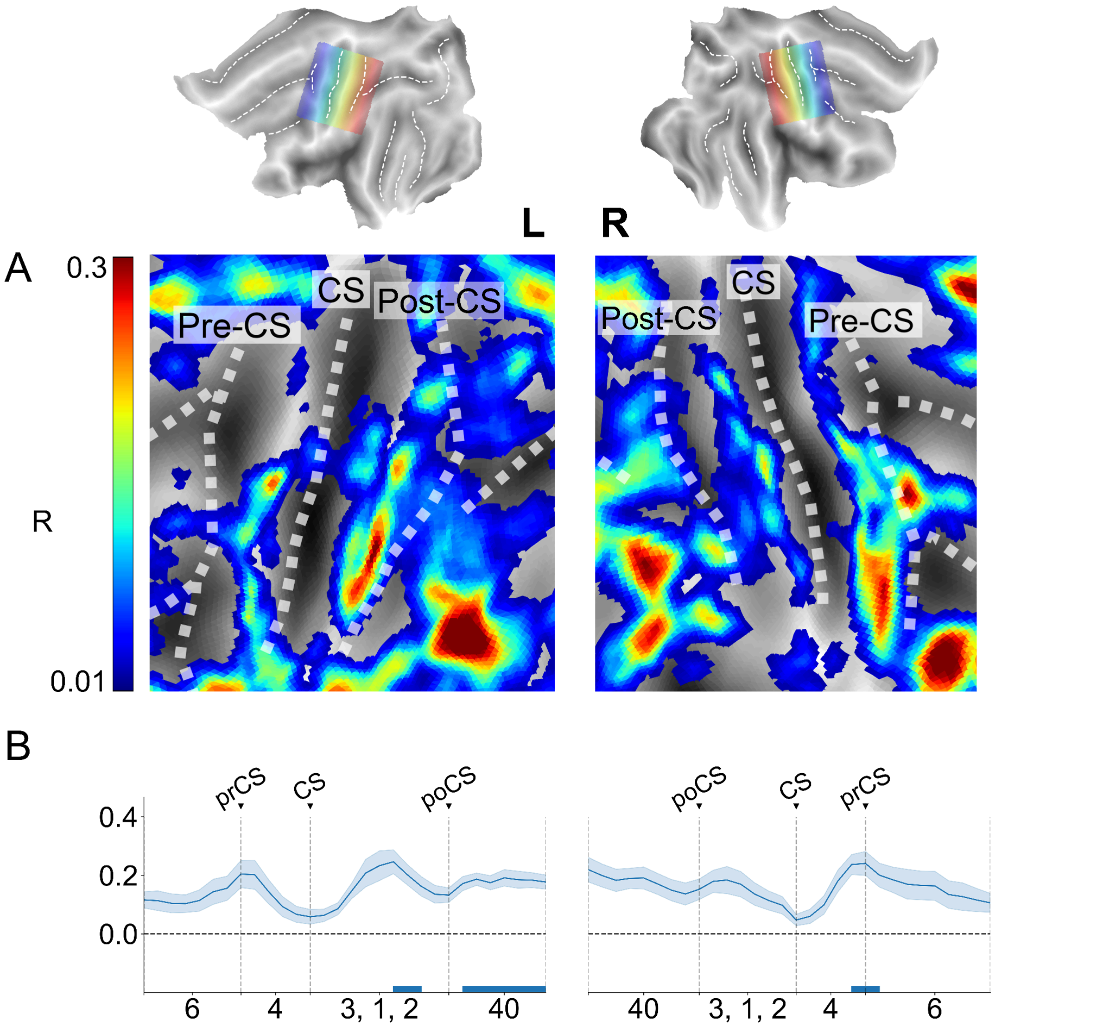


**Fig. S6: Representational dissimilarity across hemispheres for left finger stimulation and same hand response in Experiment 3. (A)** Group averaged map of mean crossnobis dissimilarity for left finger stimulation in memory condition (voxel wise uncorrected $P < 0.005$). **(B)** Cortical profile plots along the sensorimotor strip. Mean representational dissimilarity is plotted from the precentral (prCS) to the postcentral sulcus (poCS) for left finger stimulation and same hand response in Experiment 3 (group mean ± SEM). Horizontal bars indicate significant profile segments ($p < 10^{-4}$).

**Supplementary Table S1. Brain regions showing significant activation and frequency-dependent modulation during tactile stimulation in the right index finger. (Memory condition in Experiment 1)**

| | Peak (MNI) | | | Cluster level | *z* value at |
|---|---|---|---|---|---|
| | *X* | *Y* | *Z* | (p, voxels) | peak |
| **General Linear Model** | | | | | |
| **First Frequency** | | | | | |
| *R Primary Sensory Cortex* | 39 | -15 | 59 | <<0.01 / 892 | -5.275 |
| *L Secondary Sensory Cortex* | -39 | -5 | 17 | <<0.01 / 609 | 5.086 |
| *R Superior Temporal Gyrus* | 69 | -35 | 17 | <<0.01 / 343 | 4.843 |
| *L Primary Sensory Cortex* | -51 | -19 | 59 | <<0.01 / 291 | 5.060 |
| *R Posterior Insula* | 41 | -3 | 7 | <0.02 / 69 | 4.982 |
| *R Extrastriate Visual Cortex* | 27 | -65 | -3 | <0.02 / 67 | 4.644 |
| *L Posterior Middle Temporal Gyrus* | -61 | -55 | 3 | <0.02 / 62 | 4.930 |
| *R Ventral Premotor Cortex* | 65 | 7 | 19 | <0.05 / 40 | 4.013 |
| **Second Frequency** | | | | | |
| *L Secondary Sensory Cortex* | -39 | -31 | 21 | <<0.01 / 792 | 5.148 |
| *R Superior Temporal Cortex* | 67 | -35 | 21 | <<0.01 / 336 | 5.266 |
| *R Anterior Ventral Insula* | 29 | 23 | -3 | <<0.01 / 268 | 5.044 |
| *L Primary Sensory Cortex* | -51 | -17 | 55 | <<0.01 / 197 | 4.864 |
| *L Anterior Ventral Insula* | -33 | 21 | 1 | <<0.01 / 169 | 5.327 |
| *L Medial Superior Frontal Gyrus* | -5 | 11 | 55 | <<0.01 / 166 | 5.235 |
| *L Posterior Temporoparietal Junction* | -47 | -49 | 15 | <<0.01 / 155 | 5.090 |
| *R Anterior Superior Temporal Gyrus* | 59 | -11 | -5 | < 0.01 / 133 | 4.674 |
| *R Posterior Middle Frontal Gyrus* | 55 | 19 | 31 | < 0.01 / 127 | 4.414 |
| *R Secondary Sensory Cortex* | 69 | -15 | 21 | < 0.01 / 113 | 4.452 |
| *L Cerebellar Cortex* | -19 | -49 | -21 | < 0.01 / 108 | -4.434 |
| *L Primary Visual Cortex* | -21 | -99 | -15 | < 0.01 / 106 | -4.398 |
| *L Posterior Middle Frontal Gyrus* | -45 | 31 | 23 | < 0.01 / 79 | 4.175 |
| *L Extrastriate Visual Cortex* | -27 | -89 | -23 | < 0.02 / 72 | -4.533 |
| *R Posterior Middle Frontal Gyrus* | 39 | 35 | 13 | < 0.02 / 67 | 4.170 |
| *L Anterior Inferior Frontal Junction* | -51 | 13 | 29 | < 0.02 / 61 | 4.410 |
| *L Primary Visual Cortex* | -9 | -91 | 3 | < 0.03 / 57 | 4.401 |
| *L Posterior Middle Temporal Gyrus* | -55 | -67 | 9 | < 0.03 / 51 | 4.210 |
| **Linear Mixed Effect Model** | | | | | |
| **First Frequency** | | | | | |

| | | | | | |
|---|---|---|---|---|---|
| *No clusters survived* | | | | | |
| **Second Frequency** | | | | | |
| *L Cerebellar Cortex* | -15 | -59 | -17 | <<0.01 / 732 | -5.296 |
| *L Superior Parietal Lobule* | -5 | -53 | 59 | <<0.01 / 721 | -5.613 |
| *R Primary Sensory Cortex* | 49 | -25 | 63 | <<0.01 / 587 | -4.904 |
| *R Ventral Premotor Cortex* | 53 | -3 | 39 | <<0.01 / 248 | -5.740 |
| *R Superior Parietal Cortex* | 19 | -75 | 45 | <<0.01 / 185 | -6.029 |
| *L Primary Sensory Cortex* | -65 | -3 | 23 | <<0.01 / 175 | -5.161 |
| *R Inferior Parietal Cortex* | 29 | -53 | 59 | <<0.01 / 166 | -5.057 |
| *R Ventral Medial Parietal Cortex* | 15 | -39 | 55 | <<0.01 / 115 | -5.046 |
| *R Posterior Inferior Parietal Cortex* | 43 | -79 | 25 | <<0.01 / 107 | -4.882 |
| *L Dorsolateral Prefrontal Cortex* | -29 | 41 | 43 | <<0.01 / 96 | -4.650 |
| *L Inferior Parietal Cortex* | -25 | -75 | 45 | <<0.01 / 92 | -4.711 |
| *L Cerebellar Cortex* | -7 | -75 | -41 | <<0.01 / 85 | -4.647 |
| *L Posterior Angular Gyrus* | -43 | -55 | 57 | <<0.01 / 83 | -4.713 |
| *L Posterior Inferior Temporal Gyrus* | -41 | -61 | -7 | < 0.01 / 71 | -4.435 |
| *R Cerebellar Cortex* | 43 | -47 | -29 | < 0.01 / 66 | -4.877 |
| *L Secondary Visual Cortex* | -15 | -91 | -19 | < 0.01 / 66 | -4.889 |
| *R Pre-Supplementary Motor Area* | 17 | 7 | 69 | < 0.01 / 65 | -4.300 |
| *L Dorsolateral Prefrontal Cortex* | -33 | 55 | 21 | < 0.01 / 50 | -4.793 |
| *R Dorsolateral Prefrontal Cortex* | 33 | 43 | 47 | < 0.01 / 50 | -4.102 |
| *R Posterior Inferotemporal Cortex* | 43 | -75 | -11 | < 0.01 / 49 | -4.833 |
| *R Medial Temporal Lobe* | 13 | -39 | -9 | < 0.01 / 48 | -6.003 |
| *L Extrastriate Visual Cortex* | -31 | -49 | -17 | < 0.01 / 47 | -4.905 |
| *L Cerebellar Cortex* | -39 | -77 | -29 | < 0.01 / 45 | -4.424 |
| *L Caudal Middle Frontal Gyrus* | -43 | 25 | 33 | < 0.01 / 45 | -4.696 |
| *L Extrastriate Visual Cortex* | -29 | -57 | -7 | < 0.01 / 40 | -5.048 |
| *R Extrastriate Visual Cortex* | 55 | 9 | 41 | < 0.02 / 39 | -4.132 |
| *R Caudal Middle Frontal Gyrus* | 33 | -53 | -13 | < 0.02 / 37 | -4.657 |
| *L Posterior Fusiform Gyrus* | -45 | -69 | -15 | < 0.02 / 36 | -4.229 |
| *R Cerebellar Cortex* | 5 | -77 | -41 | < 0.03 / 35 | -4.360 |
| *L Hippocampus* | -39 | -15 | -23 | < 0.03 / 35 | -5.290 |
| *R Lateral Superior Temporal Gyrus* | 57 | -3 | -1 | < 0.03 / 35 | -4.652 |
| *R Posterior Anterior Cingulate Cortex* | 5 | 31 | 9 | < 0.03 / 32 | -4.502 |
| *R Lateral Superior Frontal Gyrus* | 1 | 53 | 49 | < 0.04 / 31 | -4.101 |
| *R Insula* | 37 | 21 | -5 | < 0.05 / 29 | -4.538 |

Abbreviations: L, left; MNI, Montreal Neurological Institute; R, right. Note: Voxel-wise $P < 0.001$.

**Supplementary Table S2. Brain regions showing significant activation and frequency-dependent modulation during tactile stimulation in the right index finger. (No-memory condition in Experiment 1)**

| | Peak (MNI) | | | Cluster level | *z* value at |
|---|---|---|---|---|---|
| | *X* | *Y* | *Z* | (p, voxels) | peak |
| **General Linear Model** | | | | | |
| **First Frequency** | | | | | |
| *L Secondary Sensory Cortex* | -53 | -21 | 21 | <0.01 / 397 | 4.875 |
| *L Primary Sensory Cortex* | -51 | -27 | 53 | <0.01 / 309 | 5.328 |
| *R Secondary Sensory Cortex* | 57 | -33 | 25 | <0.02 / 248 | 4.338 |
| *L Primary Visual Cortex* | -15 | -87 | 11 | <0.02 / 244 | 4.566 |
| *R Primary Sensory Cortex* | 35 | -27 | 63 | <0.05 / 116 | -3.786 |
| **Second Frequency** | | | | | |
| *R Primary Sensory Cortex* | 39 | -23 | 65 | <<0.01 / 582 | -3.169 |
| *L Medial Superior Frontal Gyrus* | -59 | -19 | 23 | <0.02 / 212 | 4.956 |
| *L Primary Sensory Cortex* | -51 | -27 | 53 | <0.02 / 210 | 4.165 |
| *L Secondary Sensory Cortex* | -51 | -19 | 19 | <0.03 / 191 | 4.611 |
| **Linear Mixed Effect Model** | | | | | |
| **First Frequency** | | | | | |
| *R Superior Parietal Cortex* | 19 | -67 | 29 | <0.01 / 152 | 4.100 |
| *L Primary Visual Cortex* | -15 | -97 | 3 | <0.01 / 118 | 4.212 |
| *L Primary Sensory Cortex* | -53 | -15 | 55 | <0.01 / 93 | 4.583 |
| **Second Frequency** | | | | | |
| *R Dorsal Premotor Cortex* | 49 | 11 | 3 | <<0.01 / 284 | -4.888 |
| *R Superior Temporal Cortex* | 59 | -45 | 23 | <0.01 / 151 | -4.467 |
| *R Inferior Frontal Cortex* | -45 | 17 | 7 | <0.01 / 134 | -4.210 |
| *L Dorsomedial Prefrontal Cortex* | -21 | 69 | 1 | <0.05 / 66 | 4.288 |
| *R Dorsolateral Prefrontal Cortex* | 31 | 61 | 9 | <0.05 / 66 | -3.813 |

Abbreviations: L, left; MNI, Montreal Neurological Institute; R, right. Note: Voxel-wise $P < 0.005$

**Supplementary Table S3. Brain regions showing significant activation and frequency-dependent modulation during tactile stimulation. (Memory condition in Experiment 2)**

| | Peak (MNI) | | | Cluster level | *z* value at |
|---|---|---|---|---|---|
| | *X* | *Y* | *Z* | (*P*, voxels) | peak |
| **General Linear Model** | | | | | |
| **First Frequency** | | | | | |
| *R Sensorimotor Network Cluster* | | | | | |
| *R Secondary Sensory Cortex* | | | | | |
| *R Primary Sensory Cortex* | | | | | |
| *R Primary Motor Cortex* | 49 | -19 | 17 | <<0.01 / 6305 | 6.823 |
| *R Insula* | | | | | |
| *R Medial Superior Temporal Gyrus* | | | | | |
| *R Ventral Premotor Cortex* | | | | | |
| *L Sensorimotor Network Cluster* | | | | | |
| *L Medial Superior Temporal Gyrus* | | | | | |
| *L Secondary Sensory Cortex* | -43 | -57 | 9 | <<0.01 / 5186 | 5.807 |
| *L Primary Motor Cortex* | | | | | |
| *L Superior Temporal Gyrus* | | | | | |
| *L Anterior Middle Temporal Gyrus* | | | | | |
| *L Extrastriate Visual Cortex* | -31 | -55 | -7 | <<0.01 / 1687 | 5.698 |
| *L Cerebellar Cortex* | | | | | |
| *R Cerebellar Cortex* | 21 | -87 | -35 | <<0.01 / 705 | 5.176 |
| *L Primary Sensory Cortex* | -43 | -21 | 51 | <<0.01 / 601 | -5.902 |
| *L Posterior Cingulate Cortex* | -1 | -45 | 37 | <<0.01 / 578 | 4.919 |
| *L Medial Superior Frontal Gyrus* | 11 | 7 | 47 | <<0.01 / 376 | 4.729 |

| | | | | | |
|---|---|---|---|---|---|
| *L Medial Parieto-Occipital Cortex* | -15 | -79 | 31 | <<0.01 / 326 | 5.071 |
| *R Medial Orbitofrontal Cortex* | 1 | 49 | -9 | <<0.01 / 249 | 4.567 |
| *R Cerebellar Cortex* | 47 | -57 | -35 | <<0.01 / 209 | 4.670 |
| *R Anterior Middle Temporal Gyrus* | 53 | -1 | -29 | < 0.01 / 166 | 4.565 |
| *L Fusiform Gyrus* | 27 | -51 | -7 | < 0.01 / 164 | 4.769 |
| *L Dorsolateral Prefrontal Cortex* | -19 | 49 | 37 | < 0.01 / 114 | 4.746 |
| *L Primary Motor Cortex* | -5 | -31 | 71 | < 0.01 / 104 | 4.753 |
| *L Anterior Ventral Insula* | -31 | 31 | 9 | < 0.02 / 79 | 4.446 |
| *R Medial Parieto-Occipital Cortex* | 17 | -77 | 31 | < 0.02 / 76 | 4.578 |
| *R Primary Visual Cortex* | 13 | -95 | -11 | < 0.02 / 73 | -4.188 |
| *R Anterior Middle Temporal Gyrus* | 49 | -45 | -15 | < 0.03 / 60 | 4.554 |
| *R Lateral Orbitofrontal Cortex* | 23 | 27 | -13 | < 0.03 / 60 | 4.532 |
| *L Lateral Orbitofrontal Cortex* | -39 | 35 | -15 | < 0.04 / 54 | 4.253 |
| *R Dorsolateral Prefrontal Cortex* | 27 | 41 | 43 | < 0.04 / 54 | 4.619 |
| *R Inferior Parietal Cortex* | 51 | -67 | 43 | < 0.04 / 53 | 5.029 |
| *L Posterior Insula* | -39 | -1 | -11 | < 0.05 / 47 | 4.357 |
| *R Hippocampus* | 21 | -5 | -23 | < 0.05 / 46 | 4.038 |
| **Second Frequency** | | | | | |
| *Sensorimotor Network Cluster*<br>*R Anterior Ventral Insula*<br>*R Primary Sensory Cortex*<br>*R Secondary Sensory Cortex*<br>*R Dorsal Premotor Cortex*<br>*R Supplementary Motor Area*<br>*L Supplementary Motor Area*<br>*R Primary Motor Cortex*<br>*L Dorsal Premotor Cortex*<br>*R Superior Parietal Cortex*<br>*R Dorsolateral Prefrontal Cortex* | 29 | 23 | -3 | <<0.01 / 11345 | 6.462 |
| *R Primary Visual Cortex*<br>*L Primary Visual Cortex*<br>*R Secondary Visual Cortex*<br>*L Secondary Visual Cortex* | 23 | -61 | 5 | <<0.01 / 2752 | 5.781 |
| *L Secondary Sensory Cortex*<br>*L Primary Sensory Cortex*<br>*L Ventral Premotor Cortex* | -53 | -19 | 17 | <<0.01 / 2358 | 5.299 |
| *L Insula*<br>*L Dorsolateral Prefrontal Cortex* | -33 | 19 | 5 | <<0.01 / 1101 | 6.166 |

| | | | | | |
|---|---|---|---|---|---|
| *L Cerebellar Cortex* | -45 | -61 | -33 | <<0.01 / 606 | 5.737 |
| *L Primary Visual Cortex* | 35 | -83 | -21 | <<0.01 / 332 | -5.325 |
| *R Cerebellar Cortex* | 25 | -65 | -51 | <<0.01 / 245 | 4.568 |
| *R Superior Parietal Lobule* | 53 | -59 | -1 | <<0.01 / 175 | 4.480 |
| *L Cerebellar Cortex* | -15 | -63 | -53 | <0.01 / 131 | 4.882 |
| *L Superior Parietal Cortex* | -17 | -43 | 71 | <0.01 / 120 | 4.474 |
| *L Posterior Inferior Temporal Gyrus* | -59 | -55 | 5 | <0.01 / 116 | 4.300 |
| *L Cerebellar Cortex* | -39 | -65 | -57 | <0.01 / 107 | 4.972 |
| *L Caudate* | -11 | 9 | -1 | <0.01 / 90 | 4.305 |
| *R Dorsal Midcingulate Cortex* | 9 | -29 | 51 | <0.02 / 70 | 4.840 |
| *R Thalamus* | 7 | -21 | 7 | <0.02 / 69 | 4.663 |
| *L Superior Parietal Cortex* | -15 | -77 | 49 | <0.03 / 55 | 4.154 |
| *R Putamen* | 21 | 9 | 3 | <0.03 / 54 | 4.361 |
| *L Medial Superior Parietal Cortex* | -5 | -37 | 61 | <0.04 / 52 | 4.222 |
| *L Caudal Anterior Cingulate Cortex* | -9 | 27 | 23 | <0.04 / 48 | 4.438 |
| *R Cerebellar Cortex* | 21 | -73 | -37 | <0.05 / 47 | 4.564 |
| *R Posterior Insula* | 29 | -5 | -3 | <0.05 / 45 | 4.936 |
| **Linear Mixed Effect Model** | | | | | |
| **First Frequency** | | | | | |
| *R Inferior Frontal Cortex* | 51 | 31 | -9 | <0.01 / 56 | -4.524 |
| | 51 | 23 | 11 | <0.01 / 52 | -4.220 |
| *L Cerebellar Cortex* | -9 | -73 | -25 | <0.03 / 31 | -4.235 |
| *R Supplementary Motor Area* | 1 | 7 | 61 | <0.04 / 29 | -4.431 |
| **Second Frequency** | | | | | |
| *L Anterior Temporal Cortex* | -35 | 11 | -33 | <<0.01 / 193 | -4.879 |
| *L Putamen* | -27 | -17 | 13 | <<0.01 / 179 | -4.894 |
| *R Medial Temporal Cortex* | 27 | -3 | -17 | <<0.01 / 97 | -4.466 |
| *R Thalamus* | 7 | -25 | 1 | <<0.01 / 84 | -5.220 |
| *L Posterior Fusiform Gyrus* | -43 | -71 | -5 | <0.01 / 44 | -4.734 |
| *L Anterior Temporal Cortex* | -49 | 13 | -31 | <0.01 / 41 | -4.913 |
| *L Anterior Superior Temporal Gyrus* | -51 | 5 | -17 | <0.01 / 40 | -4.907 |
| *L Insula* | -33 | 3 | 7 | <0.02 / 35 | -4.969 |
| *L Ventral Lateral Intraparietal Sulcus* | -31 | -53 | 57 | <0.02 / 35 | -4.277 |
| *L Hippocampus* | -27 | -9 | -21 | <0.02 / 33 | -4.717 |
| *L Dorsal Lateral Intraparietal Sulcus* | -25 | -49 | 39 | <0.02 / 33 | -4.773 |
| *L Lateral Temporal Cortex* | -47 | -23 | -5 | <0.03 / 29 | -4.215 |
| *L Dorsal Superior Temporal Gyrus* | -51 | -11 | 1 | <0.05 / 27 | -4.976 |

Abbreviations: L, left; MNI, Montreal Neurological Institute; R, right. Note: Voxel-wise $P < 0.001$
